# SILAKin: A novel high throughput SILAC and mass spectrometry-based assay to identify the substratome of kinases secreted by pathogens

**DOI:** 10.1101/2021.05.05.442720

**Authors:** Despina Smirlis, Florent Dingli, Valentin Sabatet, Aileen Roth, Uwe Knippchild, Damarys Loew, Gerald F. Späth, Najma Rachidi

**Author notes:** To whom correspondence should be addressed: Najma Rachidi. Institut Pasteur and INSERM U1201, Unité de Parasitologie Moléculaire et Signalisation, Paris, France.

## Abstract

Protein phosphorylation is one of the most important reversible post-translational modifications. It affects every cellular process including differentiation, metabolism and cell cycle. Eukaryotic protein kinases (ePK) catalyse the transfer of a phosphate from ATP onto proteins, which regulates fast changes in protein activity, structure or subcellular localisation. The systematic identification of substrates is thus crucial to characterise the functions of kinases and determine the pathways they regulate, and even more so when studying the impact of pathogens-excreted kinases on the host cell signal transduction. Several strategies and approaches have been used to identify substrates, but all show important limitations thus calling for the development of new efficient and more convenient approaches for kinase substrate identification.

Herein, we present SILAkin, a novel and easy method to identify substrates that is applicable to most kinases. It combines phosphatase treatment, pulse heating, *in vitro* kinase assay (IVKA) and SILAC (Stable Isotope Labeling with Amino acids in Cell culture)-based quantitative mass spectrometry (MS). We developed SILAkin using the *Leishmania* casein kinase 1 (L-CK1.2) as experimental model. *Leishmania*, an intracellular parasite causing Leishmaniasis, releases L-CK1.2 in its host cell. Applying this novel assay allowed us to gain unprecedented insight into host-pathogen interactions through the identification of host substrates phosphorylated by pathogen-excreted kinases. We identified 225 substrates, including 85% previously unknown that represent novel mammalian CK1 targets, and defined a novel CK1 phosphorylation motif. The substratome was validated experimentally by L-CK1.2 and human CK1δ, demonstrating the efficiency of SILAkin to identify new substrates and revealing novel regulatory pathways. Finally, SILAkin was instrumental in highlighting host pathways potentially regulated by L-CK1.2 in *Leishmania-*infected host cells, described by the GO terms ‘viral & symbiotic interaction’, ‘apoptosis’, ‘actin cytoskeleton organisation’, and ‘RNA processing and splicing’. SILAkin thus can generate important mechanistic insights into the signalling of host subversion by these parasites and other microbial pathogen adapted for intracellular survival.

## INTRODUCTION

Protein phosphorylation, one of the essential reversible post-translational modifications, affects every cellular process including transport, metabolism and DNA repair [reviewed in ^1^]. Eukaryotic protein kinases (ePK) catalyse the transfer of a phosphate from ATP onto proteins to regulate fast changes in protein activity, structure, interaction, or subcellular localisation. ePKs target proteins through kinase-specific recognition motif, which allows for specificity and restricts the number of their substrates. The systematic identification of substrates is thus crucial to characterise the functions of kinases and determine the pathways they regulate - even more so when studying the impact of pathogens-derived kinases on the host cell signal transduction during intracellular infection, for instance, by viral and bacterial pathogens ^2^. However, the low stoichiometry of protein phosphorylation, the presence of endogenous kinases as well as the reversibility of the phosphorylation by phosphatases render systematic mapping of the cellular substratome extremely challenging. This is particularly true when handling pleiotropic signalling kinases such as casein kinase 1 (CK1), able to phosphorylate hundreds of substrates ^3^, or investigating pathogen-excreted kinases that are low in abundance and compete with high-abundant host kinases thus precluding the identification of *de novo* phosphorylation events by these exo-kinases.

Several strategies and approaches have been developed and applied to identify substrates (for review see ^4^), including (i) genetic screens ^5, 6^, (ii) protein and peptide arrays ^7, 8^, (iii) phage display ^9, 10^, (iv) <u>K</u>inas<u>E</u> <u>S</u>ubstrate <u>TR</u>acking and <u>E</u>Lucidation (KESTREL) ^11^, (v) <u>K</u>inase-Interacting <u>S</u>ubstrate <u>S</u>creening (KISS)^12^, (vi) chemical-genetic methods with genetically engineered kinases ^13^ and (vii) quantitative phosphoproteomics in intact cells ^14, 15^. Although these approaches allowed the identification of an important number of substrates, but all show important limitations, which include bottlenecks inherent to high-throughput mutagenesis in genetic screens; laborious process and low-throughput screening for KESTREL; unstable interactions for KISS or the low catalytic efficiency of certain engineered kinases for chemical-genetic screens. Thus there is an urgent need to develop new easy and efficient approaches for kinase substrate identification, especially since the most important human diseases such as cancer or neurodegenerative diseases are linked to deregulation, or mutations of kinases ^16^.

Here, we pioneered a novel technology, applicable to most protein kinases, termed SILAkin that allows for easy and efficient identification of substrates. Applied to intracellular macrophage infection with the protozoan pathogen *Leishmania*, we identified host substrates that shed important new light on parasite immune subversion and validate SILAkin as a powerful new tool to study mechanisms of host/pathogen interaction.

## RESULTS

### Background

Pathogens that invade mammalian host cells export proteins and particularly kinases via exosomes to modify their host cell ^17, 18^. Identification of host substrates for these exo-kinases could be particularly challenging, as only a small fraction of the kinase pool is exported, and exosomes represent a complex environment containing many proteins, including other kinases that could mask substrate phosphorylation by the kinase of interest (KOI). We thus developed a method to identify host substrates of pathogen kinases, irrespective of their abundance and the complexity of the environment. As a proof of principle, we identified the host substrates of L-CK1.2, released by *Leishmania* parasites^18, 19^; since little is known about the pathways it regulates in macrophages. *Leishmania* is an intracellular parasite that resides in the parasitophorus vacuole of macrophages and subvert its host cell to survive.

### Strategy and establishment of the experimental protocol

To identify the L-CK1.2 host-substratome, we implemented an experimental workflow designed to quantify phospho-peptide stoichiometry by LC/MS-MS, in metabolically-labelled, heat inactivated THP-1 macrophage protein lysates after their phosphorylation by recombinant L-CK1.2 (Fig.1A). Because CK1 recognises consensus sites that require or not priming by upstream kinases, prior *in vitro* kinase assay (IVKA), we depleted the lysates in ATP before treating them or not with Antarctic phosphatase. Dephosphorylation of existing phospho-sites increased the number of sites available for *de novo* L-CK1.2 phosphorylation. Phospho-peptide stoichiometry was calculated from technical triplicate as a ratio of Heavy_L-CK1.2_/Light_L-CK1.2-K40A_ and comparisons were made to phospho-peptide ratios of Heavy/Light mock reactions without kinases.

**Figure 1:**
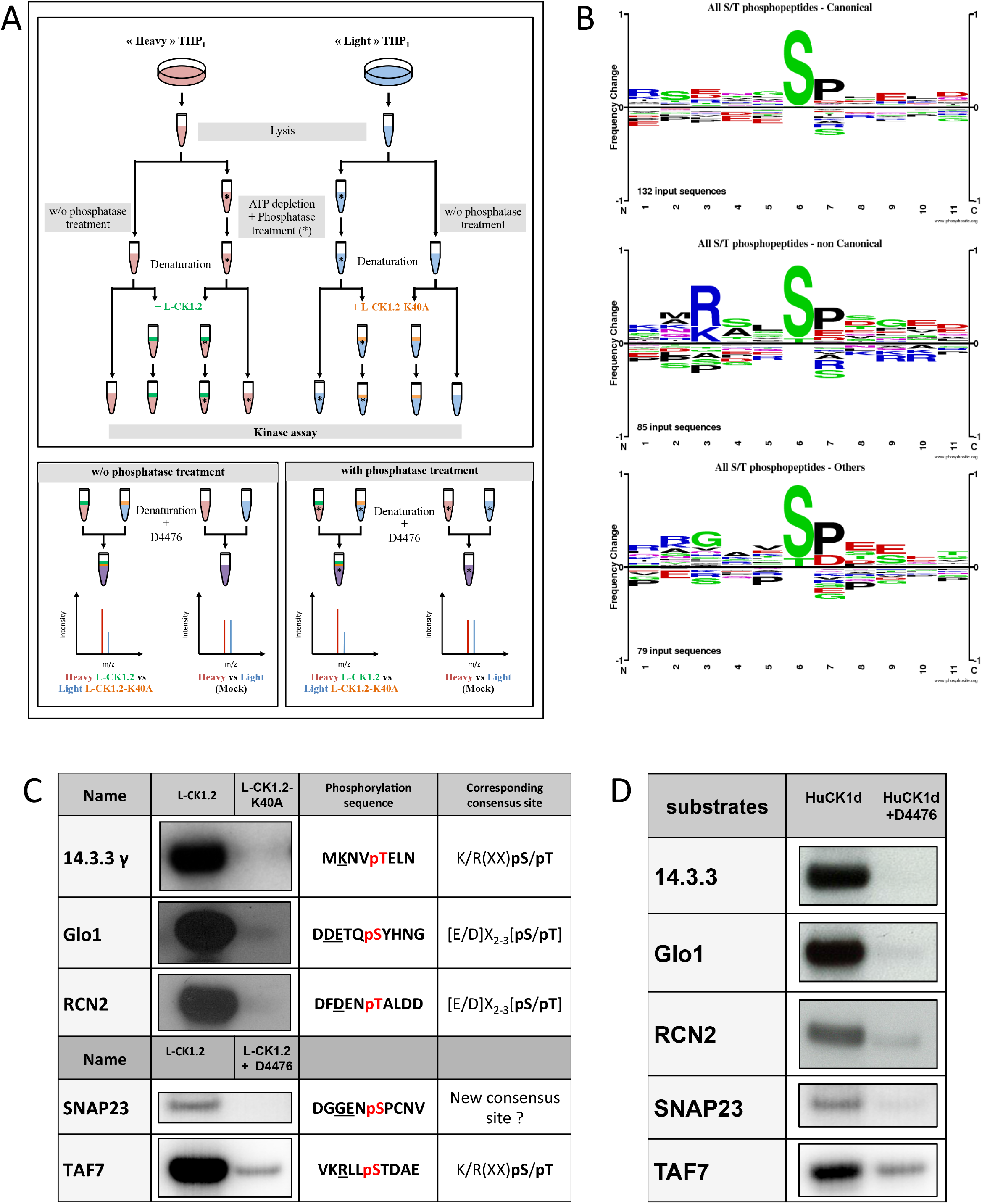
Experimental workflow and validation of SILAkin. A. Upper Panel. Workflow diagram showing the experimental strategy used to reveal L-CK1.2 substratome derived from THP-1 lysates. THP1 cells were cultured and differentiated in the presence of natural amino acids (light, blue) or stable amino acid isotopes (heavy, red). Equal amounts per reaction (0.5 mg) of heavy or light lysates were treated with phosphatase and ATP depleted (*) or not and denatured by heat inactivation to remove endogenous kinase activities. The phosphatase reactions were stopped by heat inactivation. Lysates were then subjected to IVKA in presence of recombinant L-CK1.2 (green), L-CK1.2-K40A (kinase-dead, orange), or were mock treated with equal amounts of kinase elution buffer, in triplicate. The reactions were stopped with heat inactivation and addition of 10 μM D4476. Lower panel. Equal amounts (0.5 mg) of heavy (L-CK1.2) and light (L-CK1.2-K40A) samples were mixed. In addition, mock heavy and light samples were mixed in a 1:1 ratio and used as a control. The four samples were reduced, alkylated and digested and the resulting phosphopeptides were enriched by TiO2-affinity chromatography, and processed by LC-MS/MS analysis on Orbitrap fusion. **B.** Sequence logos analysis of unique phospho-sites (five amino acids before and after the phosphorylation residues) matching strict selection criteria. Upper panel, for canonical consensus sites; middle panel, for non-canonical sites; and lower panel, for others. The amino acids are labelled according to their chemical properties: green for polar amino acids (G, S, T, Y, C, Q, N), blue for basic amino acids (K, R, H), red for acidic amino acids (D, E), and black for hydrophobic amino acids (A, V, L, I, P, W, F, M). **C.** Autoradiogram representing IVKA using selected recombinant human proteins and inactive L-CK1.2-K40A (kinase-dead), active L-CK1.2 alone or in presence of the CK1-specific small molecule inhibitor D4476 (10 μM). **D.** Autoradiogram representing IVKA using selected recombinant human proteins and recombinant GST-CK1δ^TV1^ alone or in the presence of D4476.

For establishing appropriate experimental conditions, several pilot experiments were carried out. To decrease the background activity of endogenous kinases, the lysate of THP-1 macrophages was pulse-heated to denature endogenous kinases (**Fig. S1**). Denaturation efficiency was demonstrated by the absence of ^32^P incorporation in the background control (**Fig. S1, lane 2**). In contrast, the *de novo* phosphorylation of denatured THP-1 proteins by active L-CK1.2 was detected, showing that L-CK1.2 phosphorylates proteins present in the macrophage lysate (**Fig. S1, lane 1**). To increase the number of sites available for *de novo* phosphorylation in the macrophage lysate, two steps were added to the pipeline (**Fig. 1A**). Free ATP was depleted from protein lysates by dialysis to prevent any phosphorylation while the samples were dephosphorylated by Antarctic phosphatase. The increase of ^32^P incorporation into substrates following phosphatase treatment (**Figure S1, lane 3**) confirms that previously many sites were inaccessible to *de novo* phosphorylation. To reduce technical errors during sample preparation for MS, limit missing values and perform quantitative analyses, we used Stable Isotope Labelling with Amino acids in Cell culture (SILAC). This method relies on the metabolic incorporation of either “Heavy” [^2^H_4_-Lysine (Lys^4^) and ^13^C_6_-Arginine (Arg^6^)] or natural (“Light”) [lysine (Lys^0^) and arginine (Arg^0^)] amino acids ^20, 21^. We validated by LC-MS/MS analysis the percentage of incorporation of Lys^4^ and Arg^6^ in macrophage proteins. This analysis revealed that more than 99 % of the identified peptides contained the two heavy amino acids. Finally, to reduce the risk of selecting false positive signals, we performed mock kinase assays using the “light” and “heavy” macrophage lysates without adding the kinases, to discard sites already differentially phosphorylated prior to the kinase assays.

### Application of the pipeline for the identification of L-CK1.2 host cell substrates

Three independent reactions for each condition were carried out, and two independent protein extracts treated or not with phosphatase were used. In the absence of phosphatase treatment, 7,752 unique phosphopeptides belonging to 3,544 unique proteins were identified of which 65% were quantified (see **Fig.S2A&B** for Venn diagrams). The same analysis was performed with samples pre-treated with phosphatase, and 15,852 phosphopeptides (5,158 proteins) were identified of which 66% were quantified (**Fig.S2C&D**), demonstrating the efficiency of the phosphatase treatment. However, the increase in phospho-peptides also in the H/L control (**Fig.S2A&B**) suggests that the pulse-heating step did not completely abrogate but only reduced the activity of endogenous kinases to levels undetectable by autoradiography (**Fig.S1**).

To determine *L-CK1.2* substrates, the following selection criteria were applied: (i) A probability of correct localisation of the phosphorylation site on validated peptides greater than 75%, as calculated by PtmRS software (see Data analysis of Material & methods), (ii) the ratio Heavy_L-CK1.2_/Light_L-CK1.2-K40A_ equal or above two to select the sites phosphorylated by L-CK1.2; and (iii) the ratio Heavy_L-CK1.2_/Light_L-CK1.2-K40A_ at least twofold higher than that of the control Heavy/Light to only select the newly phosphorylated sites. Using the above criteria, 99 phosphopeptides (81 proteins) and 158 phosphopeptides (144 proteins) were selected as potential substrates of *L-CK1.2* in absence or in presence of phosphatase, respectively (**Table 1**). Only fifteen proteins were common to the two datasets. Although consistent with the stochasticity of the LC/MS analysis, this finding suggests that the phosphatase treatment greatly improves the access of L-CK1.2 to new substrates for which it has more affinity.

**Table 1.** Phosphopeptides.

|  | w/o phosphatase treatment | with phosphatase treatment | Total phospho-sites | % |
| --- | --- | --- | --- | --- |
| Total phospho-peptides | 99 | 158 | 257 |  |
| **Consensus site for CK1** |  |  |  |  |
| Canonical sites | 42 | 84 | 126 | 72% |
| Non-canonical sites | 31 | 43 | 74 |  |
| Others sites | 31 | 46 | 77 | 28% |
| **Sites detected in human cell phospho-proteomes** |  |  |  |  |
| Site phosphorylated *in vivo* | 92 | 149 | 241 | 94% |
| **SARS-coV2 infection** |  |  |  |  |
| Sites phosphorylated during SARS-coV2 infection | 20 | 28 | 52 | 20% |
| **Tumorigenesis** |  |  |  |  |
| Site phosphorylated in cancer cells | 83 | 146 | 229 | 89% |
| Sites mutated during tumorigenesis | 10 | 16 | 26 | 10% |

### Validation of the SILAkin method

Three levels of validation were used to demonstrate that the dataset containing 225 proteins are bona fide host *L-CK1.2* substrates (Table 2). First, we identified 11 known human CK1 substrates, including the interferon-alpha/beta receptor alpha chain (IFNAR1) ^22^, Sprouty 2 (SPRY2) ^23^ and Fam83H ^24^ as well as 66 known CK1 binding partners, which thus might also be substrates. Second, we showed that the phosphorylated peptides were highly enriched in known CK1 consensus sites^25^. Hundred and twenty-six sites display the canonical CK1-phosphorylation motif, [S/T]X_2-3_[pS/pT] or [E/D]X_2-3_[pS/pT] (Table 1, Fig. 1B top panel), consistent with the affinity of CK1 for these two motifs ^26–33^. Seventy-four sites display the non-canonical consensus SLS-Xn-(E/D)n ^29, 34–36^ or resemble to K/R(X)K/R(XX)**pS**/**pT**, a CK1 consensus site identified in cholesterol and sulfatide binding proteins (**Table 1, Fig. 1B middle panel**)^37^. Surprisingly, this motif was present in only 10 phosphopeptides while the remaining 58 peptides contained a shorter version, K/R(XX)**pS**/**pT**. Finally, seventy-seven phospho-sites did not contain any known CK1 motif (others, **Table 1**), and might represent novel mammalian CK1 phosphorylation motif (**Table 1**). Indeed, we identified [G]XX[**pS**/**pT**] in 13 phospho-peptides (**Fig. 1B bottom panel and Table 3**). Noticeably, all the consensus sites identified in this study have a proline residue adjacent to the phosphorylated S or T, [**pS**/**pT**][P] (**Fig. 1B, position 7**), which was not described for other CK1s. In that, it resembles the phosphorylation motif phosphorylated by CDKs^38^, and might be a characteristic of L-CK12 substrate recognition. Third, experimental validation of the phosphoproteomics analyses was carried out by the implementation of an *in vitro* kinase assay (IVKA) using purified human recombinant proteins, as substrates. We included five proteins that might be important for *Leishmania* intracellular survival based on their potential to modulate macrophage functions and/or inflammation. The selected recombinant proteins, namely reticulocalbin 2 (RCN2, Q14257), 14-3-3γ (YWHAG, P61981), lactoylglutathione lyase 1 (Glo1, Q04760), synaptosomal-associated prot 23 (SNAP23, O00161) and transcription initiation factor TFIID subunit 7 (TAF7, Q15545) were subjected to a kinase assay using inactive L-CK1.2-K40A (kinase-dead), active L-CK1.2 alone or in presence of the CK1-specific small molecule inhibitor D4476 ^39^. All recombinant proteins were phosphorylated by L-CK1.2, but no phosphorylation was observed in the D4476 and L-CK1.2-K40A controls (**Fig. 1C**). All L-CK1.2 substrates tested were also phosphorylated by CK1δ, confirming the relevance of our substratome for human CK1s (**Fig. 1D**). Altogether, these results confirm that our dataset identified bona fide L-CK1.2 host cell substrates, which validate the SILAkin method. Furthermore these substrates are potentially also targeted by human CK1δ, revealing novel targets and thus new pathways regulated by human CK1δ.

**Table 2.**
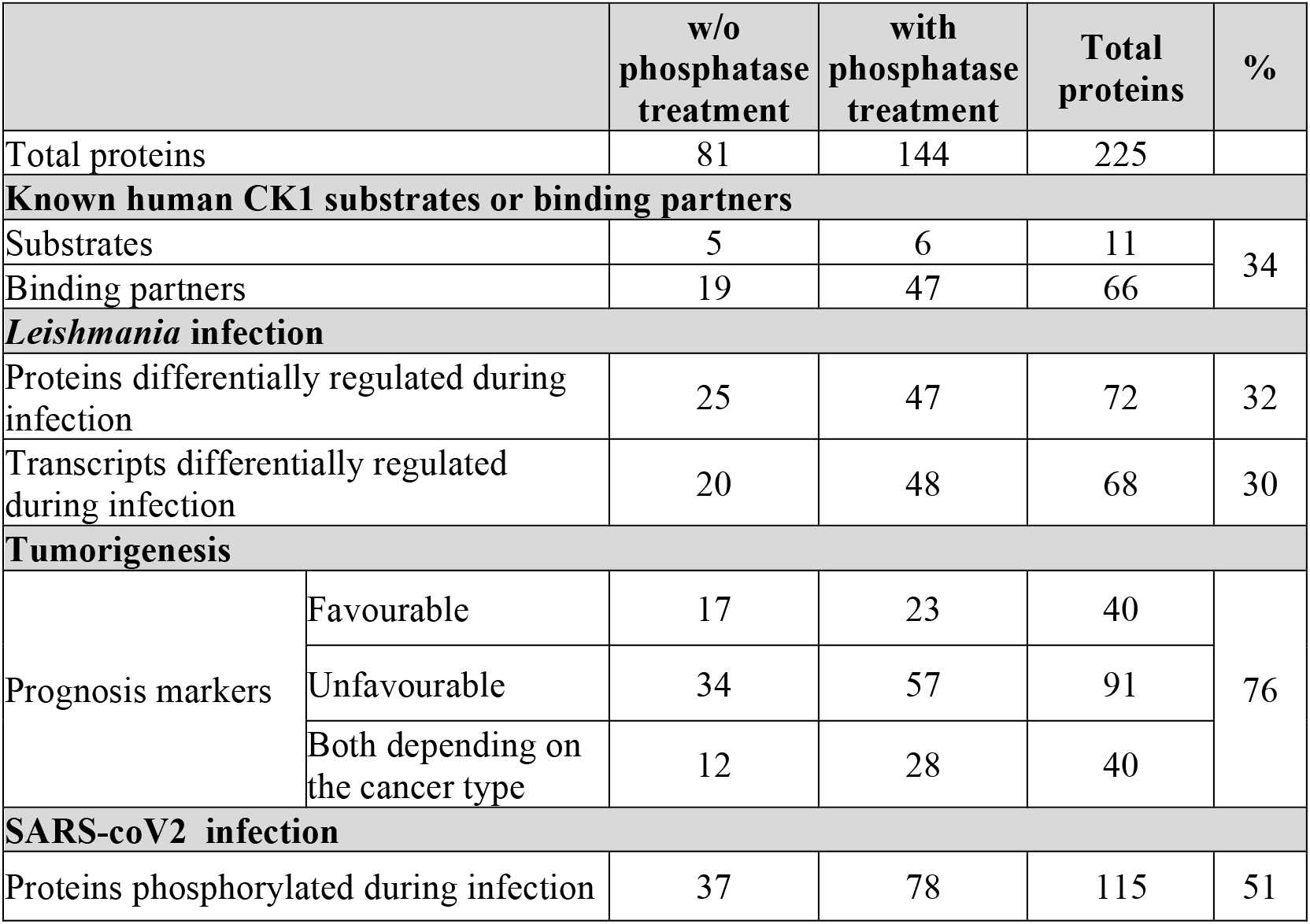
Proteins.

**Table 3.** Motifs identified in category ‘others’.

| Motif | ZScore | P-Value | Foreground Matches | Foreground Size |
| --- | --- | --- | --- | --- |
| ..G..sP.... | 8.64 | 0.00E+00 | 10 | 69 |
| ..G..s..E.. | 8.24 | 0.00E+00 | 7 | 69 |
| ..G..s..... | 6.4 | 8.02E-11 | 20 | 69 |
| .....sP.... | 9.94 | 0.00E+00 | 34 | 69 |
| .....sD.E.. | 8.43 | 0.00E+00 | 8 | 69 |
| .....sD.... | 8.73 | 0.00E+00 | 11 | 69 |
| .....s..E.. | 5.03 | 2.43E-07 | 16 | 69 |

### Physiological relevance of L-CK1.2 substrates for human diseases

Because SILAkin is an *in vitro* method, the question of physiological relevance needed to be addressed. To this end, our dataset was compared to human cell phospho-proteomes to determine whether any of the 257 phospho-sites were phosphorylated *in vivo*. 94% were identified in various phospho-proteomes (Table 1 and S1, https://www.phosphosite.org/homeAction.action), suggesting that the phospho-sites identified by SILAkin are physiologically relevant. Our data allow the attribution of those sites to human CK1. Similarly, we investigated whether any L-CK1.2 substrates were described as differentially regulated in human diseases. Indeed, 89% of phospho-sites are phosphorylated in cancer cells (**Table 1&S1 and Fig.2, round shape**) and 10% are mutated during tumorigenesis (**Table 1&S1 and Fig.2, red border**). Moreover, 76% of the L-CK1.2 substrates are considered as prognosis markers for various cancer types (**Table 1&S1**). These data are consistent with the fact that human CK1 isoforms are overexpressed in cancer cells ^25, 40^ and often contribute to chemo- and apoptosis resistance as well as to a reduction of overall survival of patients ^25^. Regarding infectious diseases, we showed that 51% of L-CK1.2 substrates are phosphorylated during SARSCoV2 infection (**Fig. 2, green**), and 20% on the same sites as those phosphorylated by L-CK1.2 (**Table 1&S1**). Surprisingly, only two GO processes are enriched, ‘RNA splicing’ and ‘cellular localisation and transport’ in the substrates phosphorylated by L-CK1.2 and SARS-CoV2 infection (**Fig. S3 and Table S2&S3**), suggesting that SARS-CoV2 and *Leishmania* might similarly exploit these two processes and that SARS-CoV2 might exploit CK1 pathways for its replication. Our data demonstrate the potential impact of CK1 in SARS-CoV2 infection, which is consistent with the role of host CK1 in viral replication ^41^ ^42–44^.

**Figure 2.**
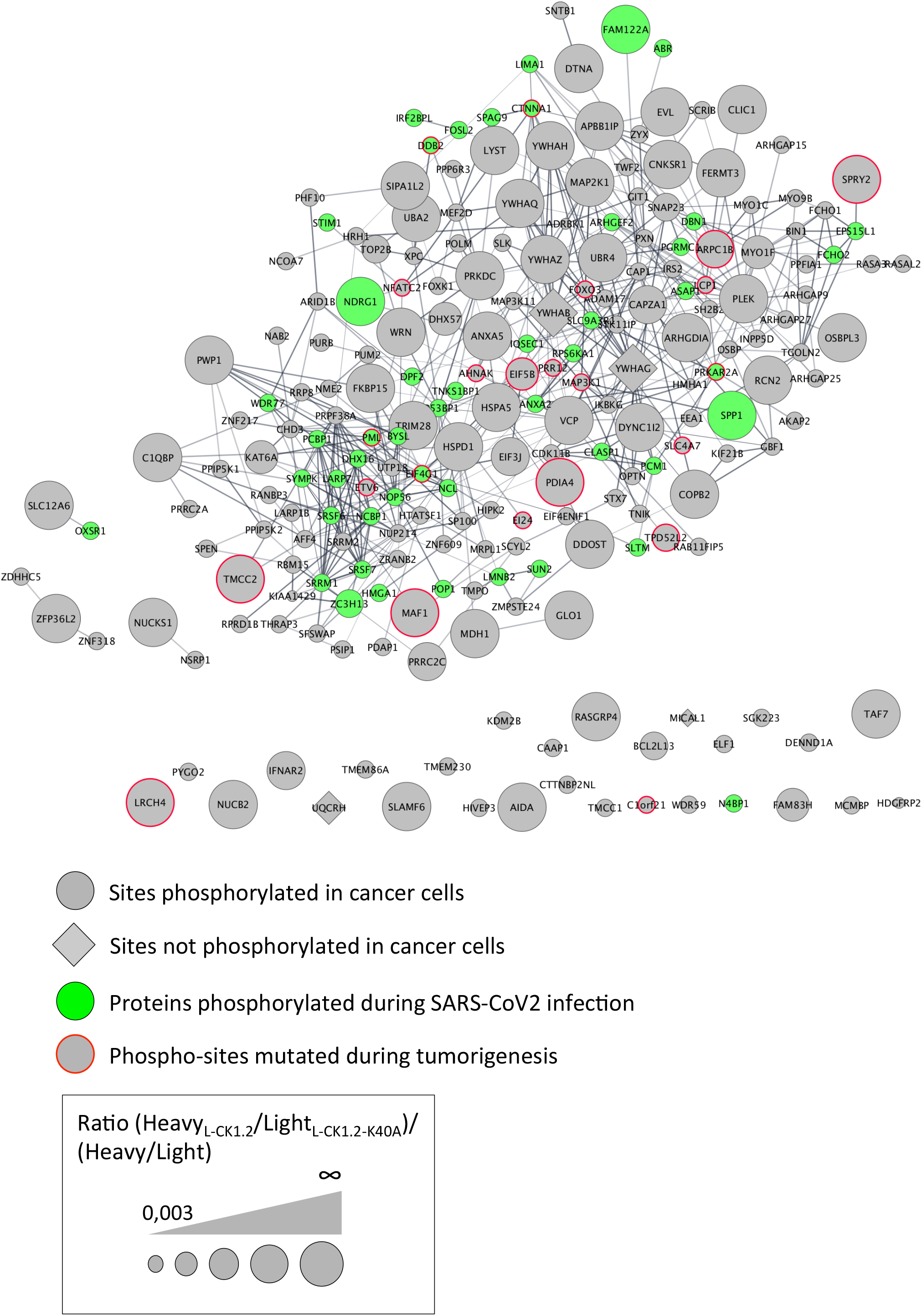
Protein-protein interaction network of L-CK1.2 substrates. The dataset was analyzed for protein-protein interactions and visualized using the STRING plugin of the Cytoscape software package. Each node represents a substrate and each edge represents a proteinprotein interaction. Round shape represents sites phosphorylated in cancer cells; Diamond shape represents sites not phosphorylated in cancer cells; Red border indicates phospho-sites mutated during tumorigenesis; Green fill color indicates proteins phosphorylated during SARS-Cov2 infection. The labeling indicates the UniProt human name. The size of the node represents the ratio (Heavy_L-CK1.2_/Light_L-CK1.2-K40A_)/(Heavy/Light mock).

### Enriched pathways and relevance for Leishmania infection

Although obtained *in vitro*, the substratome is enriched in proteins physiologically relevant for *Leishmania* infection (60% total) and might give us important insights into the mechanisms developed by *Leishmania* to subvert its host cell. Indeed, during *Leishmania* infection, 32% of L-CK1.2 substrates were shown to be differentially regulated at protein level ^45–48^, while 30% were differentially expressed at transcript level ^49–52^ (Table 1). In the substratome, we identified functional enrichment for GO terms relative to apoptosis or RNA processing and splicing, which is consistent with *Leishmania* inhibiting host apoptosis ^53, 54^ or modifying the host transcriptome, respectively ^49, 55^ (**Fig. 3**). Moreover, several key processes, preferentially targeted by *Leishmania* CK1.2, are associated with host-pathogen interactions such as ‘viral and symbiotic interaction’ or ‘cellular response stimulus’ (**Fig. 3, Table S4**). Lastly, the identification of processes such as ‘positive regulation of metabolic process’, ‘positive regulation of molecular function activity’ and ‘GTPase signal transduction’, are consistent with L-CK1.2 being a signalling kinase (**Fig. 3, Table S4**). Altogether, its functional characterisation supports the physiological relevance of the substratome, while revealing the pathways that might be regulated by L-CK1.2 during *Leishmania* infection. These pathways should be investigated in priority to decipher L-CK1.2 functions in the host cell.

**Figure 3.**
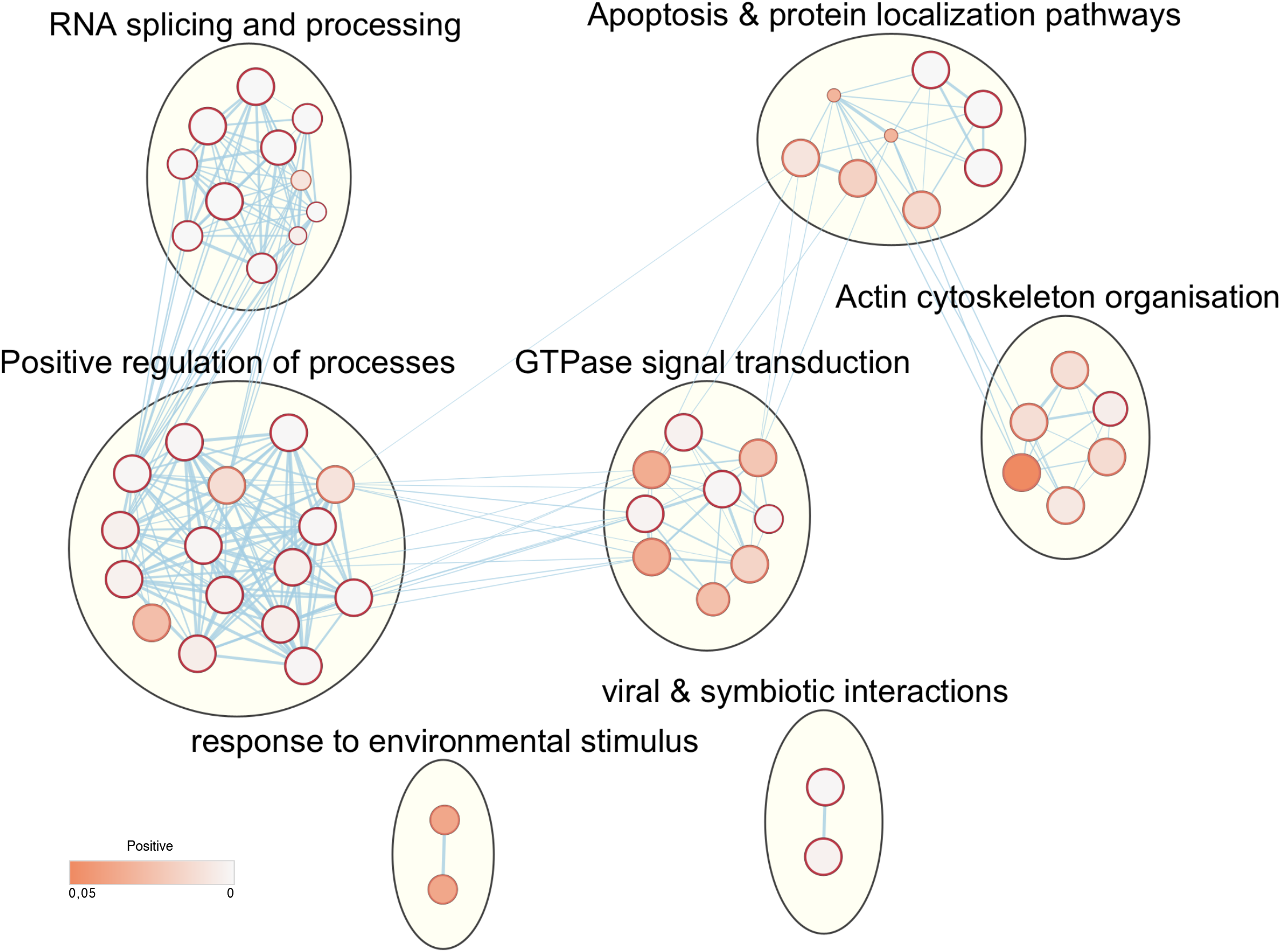
L-CK1.2 targets specific pathways. Functional enrichment analysis of the whole dataset was performed using the g:profiler web server. Results were visualized using the EnrichmentMap plugin of the Cytoscape software package, with a pvalue and a Q-value above 0.05 and an edge cut-off of 0.375. Node colour represents the enrichment p-value. Node size is proportional to the total number of genes belonging to the corresponding geneset. The edge corresponds to the Annotation shared between two nodes (blue), with edge thickness corresponding to the number of shared genes. Node clusters were identified and annotated by the AutoAnnotate plugin of cytoscape. See Table S1 for list of the whole list of annotations.

## DISCUSSION

Increasing the knowledge on kinases and particularly the pathways they regulate is crucial to better understand cellular functions, hence the importance of finding their substrates in an unbiased manner. It is even more important when studying host cell signalling pathways exploited by pathogens during infection through the release of their kinases. Many high throughput methods exist and most of them have important drawbacks or are not adapted to host-pathogen interaction studies ^56^. Here, we propose SILAkin, a novel, easy, and validated method applicable to most kinases from any organism that overcomes most of the limitations existing in other mass spectrometry-based methods: First, it allows direct detection of phosphorylation, contrary to phospho-proteomics comparing WT and mutant kinase ^4^. Second, it does not require the modification of the kinase, thus avoiding potential artifacts observed in chemical-genetic methods ^13^. Finally, ATP depletion and phosphatase treatment were particularly efficient to reveal novel and previously inaccessible phospho-sites. Our method has two main limitations. First, the variability of mass spectrometry analyses, thus increasing the number of replicate will increase the depth of identification and consequently the number of substrate identified. Second, limitations inherent to *In vitro* kinase assays such as non-physiological kinase regulation, sub-optimal production of an active recombinant kinase or *in vivo* substrate accessibility, thus *in vivo* validation of the dataset will be required. Because L-CK1.2 is constitutively active, accesses all cellular compartments ^39, 57^, and recognises specific consensus sites, the limitations inherent to IVKA do not apply. L-CK1.2 is therefore the perfect kinase to demonstrate the efficiency of SILAkin.

Our large dataset allowed us to refine existing CK1 substrate recognition motif such as K/R(X)K/R(XX)**pS**/**pT**, for which we proposed a shorter version, K/R(XX)**pS**/**pT** ^37^; and to define a novel CK1 consensus site such as [G]X_2-3_[**pS**/**pT**], validated experimentally with L-CK1.2 and CK1δ (SNAP23, **Fig. 1C and D**). Further analyses, including mutagenesis, will be required to confirm this new consensus, as we cannot exclude the possibility that SNAP23 was phosphorylated on another site. Surprisingly, we identified a proline residue adjacent to the phosphorylated S/T ([**pS**/**pT**][P]), its requirement remains to be confirmed experimentally and might only applied to *Leishmania* CK1.2, since it was not described for other CK1 orthologs.

Finally, this study identified 225 CK1 substrates, including 214 previously unknown. Many substrates have a link with human diseases, including cancer, SARS-CoV2 ^58^ or *Leishmania* infections ^46–48, 59^, suggesting that the corresponding pathways might be regulated by CK1. These findings are consistent with human CK1 having cancer-associated functions linked to its involvement in the Wnt (Wingless/Int-1), Hh (Hedgehog), and Hippo signalling pathways ^3, 25^, or with the involvement of CK1 in infectious diseases ^60–63^. Applied to *Leishmania* parasites, this study by revealing the potential pathways regulated by L-CK1.2 in macrophages, such as ‘viral & symbiotic interaction’, ‘actin cytoskeleton organisation’ ^64^, and ‘RNA processing and splicing’ ^49, 55^, proposes mechanistic insights into the signalling of host subversion by *Leishmania*.

In conclusion, we developed a method sufficiently sensitive to reveal substrates phosphorylated by pathogen-excreted kinases, allowing us to gain insights into host-pathogen interactions.

## MATERIALS & METHODS

### SILAC labelling and lysate preparation

For labelling cells by SILAC, equal numbers of THP-1 monocytes (2*10^5^ ml^−1^) were seeded in RPMI 1640 without Lysine and Arginine (Thermo Fisher Scientific), supplemented either with natural amino acids (L-Lysine, 0.274 mM; L-Lysine, 1.15 mM; Arginine, 1.15 mM) or with the same concentrations of amino acid isotopes 2H4-Lysine (Lys4) and 13C6--Arginine (Arg6) (Thermo Scientific). The medium was supplemented with 50 μM β-mercaptoethanol, 50U mL^−1^ penicillin, 50 μgmL^−1^ streptomycin and 10% (v/v) of dialysed Fetal Bovine Serum (Sigma). Cells were split and seeded before reaching a concentration of 10^6^ mL^−1^ in fully supplemented SILAC medium, for a period of at least 15 days. 0.75 to 1*10^8^ cells were then differentiated for 48 h into macrophages by the addition of 10 ng mL^−1^ PMA. Cells cultivated in SILAC medium were washed three times in PBS and lysed in RIPA lysis and extraction buffer (Thermo Scientific) containing one tablet per 10 mL of cOmplete™ Protease Inhibitor Cocktail tablets (Sigma). Cell extracts were incubated on ice 30 min, sonicated 5 min, and centrifuged 15 min at 14,000 g to eliminate cell debris. Proteins were quantified in the supernatants using the RC DC™ protein assay kit (Bio-Rad), according to the manufacturer’s instructions. For free ATP depletion, protein extracts were dialyzed overnight at 4°C in 1 liter of dialysis solution (1× PBS, 1 mM EDTA, 1 mM dithiothreitol) in a Slide-A-Lyzer dialysis cassette (Pierce).

### Expression and purification of L-CK1.2 and L-CK1.2-K40A

Bacterial expression plasmid for L-CK1.2 was generated as previously described ^62^. Bacterial expression plasmid carrying L-CK1.2-K40A (kinase dead) was generated by site directed mutagenesis as previously reported ^62^. Recombinant proteins were induced in Rosetta™ (DE3) Competent Cells (Novagen) with 0.02% (w/v) L-arabinose for 3 h at 25 °C. Cells were harvested and resuspended in lysis buffer as previously described ^62^. Briefly recombinant kinases were purified on co-nitrilotriacetic acid agarose (Pierce) and eluted in 300 mM imidazole in PBS containing 60 mM β-glycerophosphate, 1 mM sodium vanadate, 1 mM sodium fluoride and 1 mM disodium phenylphosphate. Protein eluates were supplemented with 15% glycerol and stored at −80°C.

### Protein kinase assay and phosphatase treatment

For phosphatase treatment, 500 μg of “heavy” or “light” THP-1 protein extract were dephosphorylated with 50 U of Antarctic phosphatase (NEB) in Antarctic phosphatase buffer (NEB) for 30 min at 37 °C. Phosphatase activity was heat inactivated at 65°C for 15 min. The kinase assay for the “heavy” or “light” protein extracts (500 μg) were performed in buffer C (60 mM β-glycerophosphate, 30 mM *p*-nitrophenyl phosphate, 25 mM MOPS [morpholinepropanesulfonic acid], 5 mM EGTA, 15 mM MgCl_2_, 1 mM dithiothreitol, 0.1 mM sodium vanadate; pH 7.0) in the presence of 15 μM ATP and 200 ng of recombinant L-CK1.2 or kinase dead L-CK1.2-K40A. The reaction was performed in triplicate at 30°C for 45 min. For mock reactions no kinases were added. Reactions were stopped with the addition of 10 μM of D4476 {4- [4-(2,3-dihydro-1,4- benzodioxin-6-yl)- 5- 2-pyridinyl)- 1*H*- imidazol-2-yl]} benzamide, a specific inhibitor of CK1 known to inhibit *L-CK1.2* activity ^65^, followed by heat inactivation at 65°C for 15 min. Next, kinase inactivation heavy and light protein samples were mixed in a 1:1 ratio, and precipitated in 80 (v/v) % ice-cold acetone and stored at −80°C prior to LC/MS-MS analysis. For IVKA with L-CK1.2, 15 μg of total THP-1 protein extracts or 0.5 to 1.0 μg of recombinant proteins, GST-14-3-3γ (Enzo #BMLSE313-0100), RCN2-6XHis (Abcam #ab125644) Glo1 (Abcam # ab206792), SNAP23 (Abnova #H00008773-P01) and TAF7 (Abnova #H00006879-P01), were assayed with 0.2 μg of L-CK1.2 or L-CK1.2-K40A, as described in Rachidi *et al.* ^39^. Incorporated ^32^P was monitored by SDS-PAGE and autoradiography. For IVKA with human GST-CK1δ^TV1^, 6 ng of kinase was incubated with the 0.5 to 1.0 μg of recombinant proteins (see above) in 1x kinase buffer (250mM tris-HCL pH 7.0, 100 mM MgCl_2,_ 1mM EDTA) and 15μM ^32^P-γATP in 30μL final volume for 30 min at 30°C. All kinase assays were performed at least three times.

### Phospho-peptide enrichment

Phosphorylated peptides were enriched using Titansphere^™^ Phos-TiO kit centrifuge columns (3mg/200μL, cat. No. 5010-21312, GL Sciences), as described by the manufacturer. After elution from the Spin tips, phospho-peptides were vacuum concentrated to dryness and reconstituted in 0.1% formic acid. Samples were then loaded onto a custom-made C18 StageTips for desalting. Enriched phosphor-peptides were eluted using 40/60 MeCN/H2O + 0.1% formic acid and vacuum concentrated to dryness.

### Liquid chromatography-tandem mass spectrometry

LC was performed with an RSLC nano system (Ultimate 3000, Thermo Scientific) coupled online to an Orbitrap Fusion Tribrid mass spectrometer (Thermo Scientific). Peptides were first trapped on a C18 column (75 μm inner diameter × 2 cm; nanoViper Acclaim PepMap^™^ 100, Thermo Scientific) with buffer A (2/98 MeCN/H2O in 0.1% formic acid) at a flow rate of 2.5 μL/min over 4 min. Separation was then performed on a 50 cm × 75 μm C18 column (nanoViper Acclaim PepMap^™^ RSLC, 2 μm, 100Å, Thermo Scientific) regulated to a temperature of 55°C with a linear gradient of 2% to 25% buffer B (100% MeCN in 0.1% formic acid) at a flow rate of 350 nL/min over 211 min. Separation of phospho-peptide samples without phosphatase, was performed with buffer A’ (5/98 DMSO/H2O in 0.1% formic acid) and B’ (5/95 DMSO/MeCN in 0.1% formic acid) by using a linear gradient of 2% to 30% with the same time and flow rate as previous gradient. Full-scan MS was acquired in the Orbitrap analyzer with a resolution set to 240,000, a mass range of m/z 400-1500 and a 4 × 10^5^ ion count target. Ions from each full scan were HCD fragmented with normalized collision energy of 30 % and rapid scan MS analysed in the linear ion trap. The MS^2^ ion count target was set to 2 × 10^4^ and only those precursors with charge state from 2 to 7 were sampled for MS^2^ acquisition

### Data analysis

For identification, the data were searched against the Homo sapiens (UP000005640) UniProt database using Sequest-HT through Proteome Discoverer (PD, version 2.4). Enzyme specificity was set to trypsin and a maximum of two-missed cleavage sites was allowed. Oxidized methionine, Met-loss, Met-loss-Acetyl, Ser/Thr/Tyr phosphorylation, N-terminal acetylation, heavy ^2^H_4_-Lysine (Lys4) and ^13^C_6_-Arginine (Arg6) were set as variable modifications. Carbamidomethyl of Cysteines were set as fixed modification. Maximum allowed mass deviation was set to 10 ppm for monoisotopic precursor ions and 0.6 Da for MS/MS peaks. The resulting files were further processed using myProMS v3.9.2 ^66^. The Sequest-HT target and decoy search result were validated at 1% false discovery rate (FDR) with Percolator at the peptide level. Technical replicates (n=3) were merged using the MSF files node and a SILAC-based phospho-peptides quantification was performed by computing peptides XICs (Extracted Ion Chromatograms). The phospho-site localization accuracy was estimated by using the PtmRS node in PD (version 2.4), in PhosphoRS mode only. Phospho-sites with a localization site probability greater than 75% and with at least two SILAC measurements per peptide were quantified at the peptide level. Mass spectrometry proteomics data have been deposited to the ProteomeXchange Consortium via the PRIDE ^67^ partner repository with the dataset identifier PXD.

### Motif discovery

Unique phospho-peptides sequences, matching the strict selection criteria, were aligned on phosphorylated serine and threonine with 5 flanking amino acids. PhosphoSitePlus phosphosite plus (https://www.phosphosite.org/homeAction.action^68^), was used to compute motif analysis enrichment (automatic background selection; significance of 1e-6; support threshold of 0.1) and to generate corresponding sequence logo (automatic background selection; frequency change algorithm ^69^.

### STRING network visualization and gene ontology enrichment analysis

The dataset was analyzed for protein-protein interactions and visualized using the STRING plugin (string, https://string-db.org/^70^) of the Cytoscape software package (version 3.8.2, https://cytoscape.org/^71^). Each node represents a substrate and each edge represents a protein-protein interaction. Functional enrichment analysis of the dataset was performed using the g:profiler web server (https://biit.cs.ut.ee/gprofiler/gost) using the following criteria: only annotated genes, with a significance threshold of 0.05, select GO terms of less than 5 000 genes, and only focusing on GO ‘biological process’. Results were visualized using the EnrichmentMap plugin of the Cytoscape software package (version 3.3, http://apps.cytoscape.org/apps/enrichmentmap ^72^), with a p-value and a Q-value above 0.05 and an edge cut-off of 0.375. Node color represents the enrichment p-value (see legend Figure 2). Node size is proportional to the total number of genes belonging to the corresponding gene-set. The edge corresponds to the Annotation shared between two nodes, with edge thickness corresponding to the number of shared genes. Node clusters were identified and annotated by the AutoAnnotate plugin of cytoscape (version 1.3.3 https://autoannotate.readthedocs.io/en/latest/).

## SUPPLEMENTARY DATA

**Figure S1.**
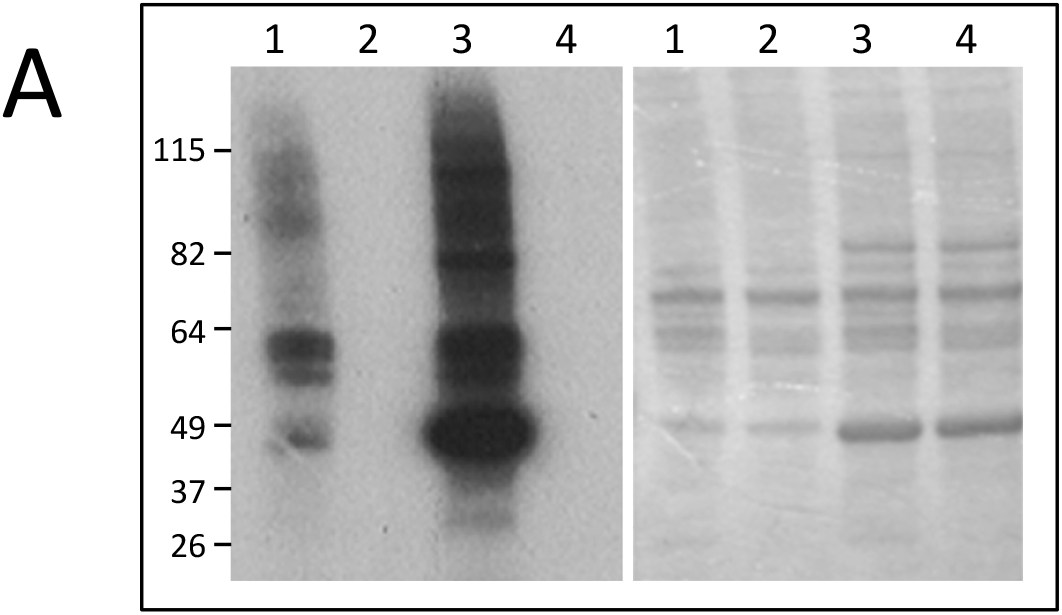
Increase of accessible residues for *de novo* phosphorylation following phosphatase treatment. Left panel. Autoradiogram representing IVKA using THP-1 cell extracts treated (lane 3 and 4) or not (lane 1 and 2) with Antarctic phosphatase, and active L-CK1.2 (lane 1 and 3) or inactive L-CK1.2-K40A (lane 2 and 4). Right Panel. Same gel stained with coomassie blue.

**Figure S2.**
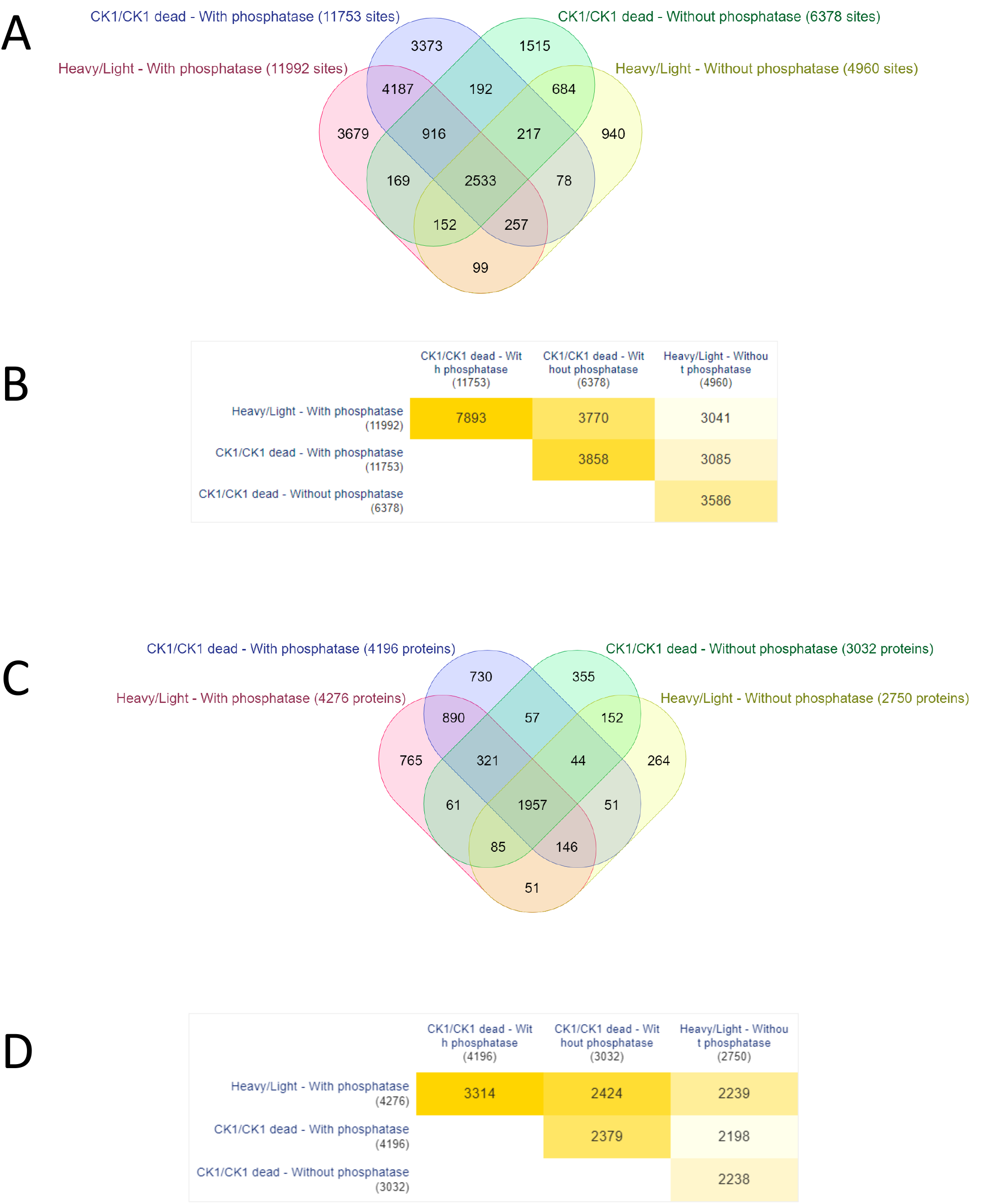
Comparison of the different experimental conditions. Venn diagrams showing the numbers and overlaps of phosphopeptides (A and C) or proteins (B and D) in the following four experiments: Heavy_L-CK1.2_/Light_L-CK1.2-K40A_ and Heavy/Light mock, treated (A and B) or not with Antarctic phosphatase (C and D).

**Figure S3.**
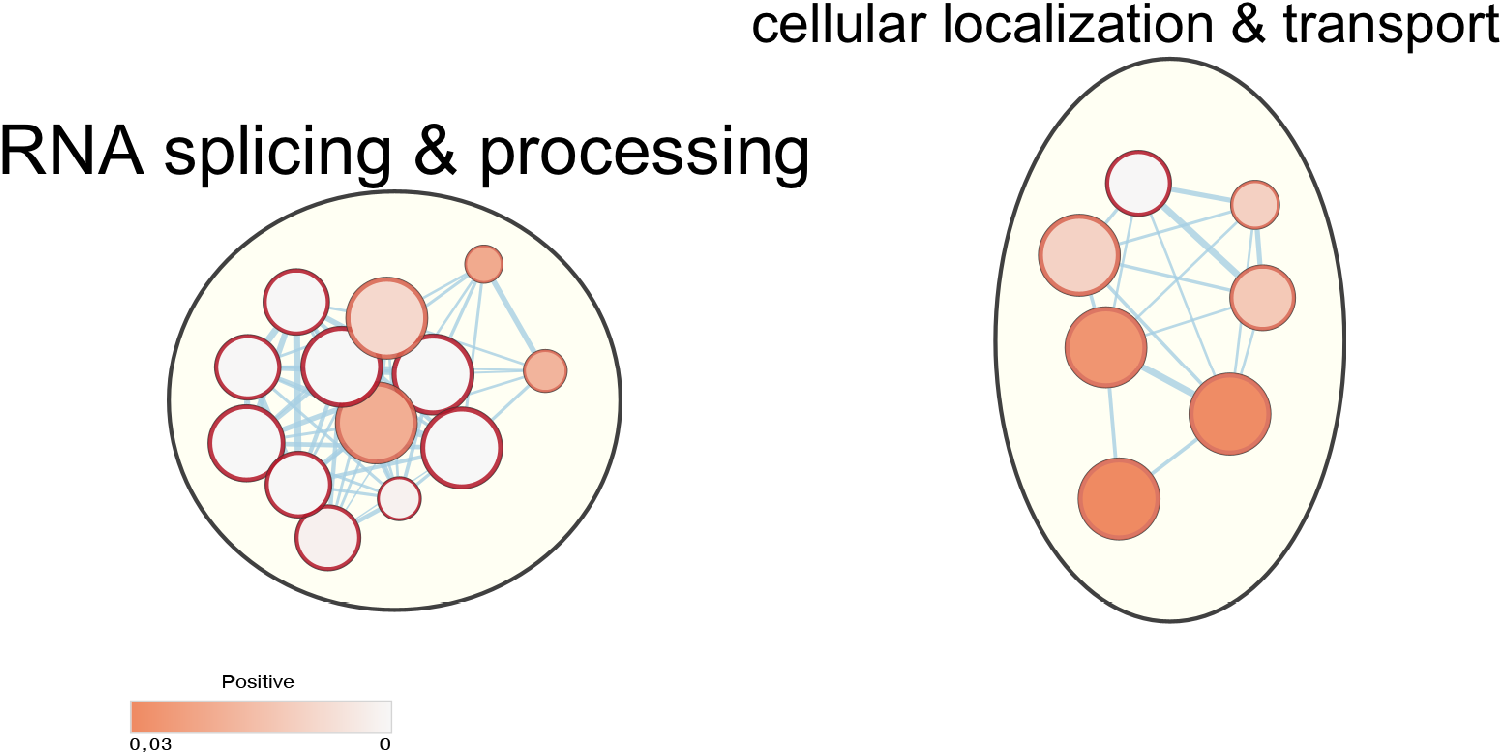
Common pathways targeted by L-CK1.2 and phosphorylated during SARS-CoV2 infection. Functional enrichment analysis of the proteins phosphorylated by L-CK1.2 and also during SARSCoV2 infection was performed using the g:profiler web server. Results were visualized using the EnrichmentMap plugin of the Cytoscape software package, with a p-value and a Q-value above 0.03 and an edge cut-off of 0.375. Node colour represents the enrichment p-value. Node size is proportional to the total number of genes belonging to the corresponding gene-set. The edge corresponds to the Annotation shared between two nodes (blue), with edge thickness corresponding to the number of shared genes. Node clusters were identified and annotated by the AutoAnnotate plugin of cytoscape. See Table 2 (RNA splicing & processing) and 3 (cellular localisation & transport) for the list of annotations.

**Table S2.** List of annota1ons of proteins targeted by LCK1.2 and phosphorylated in SARS-CoV2 infection.

| RNA splicing & processing |  |  |
| --- | --- | --- |
| Gene set name | Gene set description | fdr-qvalue |
| GO:0008380 | RNA splicing | 7.56E-06 |
| GO:1903311 | regulation of mRNA metabolic process | 0.002061234 |
| GO:0000398 | mRNA splicing, via spliceosome | 2.89E-04 |
| GO:0090304 | nucleic acid metabolic process | 0.023112918 |
| GO:0031123 | RNA 3'-end processing | 0.021130162 |
| GO:0016071 | mRNA metabolic process | 1.13E-06 |
| GO:0000375 | RNA splicing, via transesterification reactions | 3.16E-04 |
| GO:0006397 | mRNA processing | 9.25E-07 |
| GO:0050684 | regulation of mRNA processing | 0.001816144 |
| GO:0031124 | mRNA 3'-end processing | 0.024329926 |
| GO:0006396 | RNA processing | 1.05E-06 |
| GO:0000377 | RNA splicing, via transesterification reactions with bulged adenosine as nucleophile | 2.89E-04 |
| GO:0016070 | RNA metabolic process | 0.009712339 |

**Table S3.** List of annota1ons of proteins targeted by LCK1.2 and phosphorylated in SARS-CoV2 infection.

| Cellular localisation & transport |  |  |
| --- | --- | --- |
| Gene set name | Gene set description | fdr-qvalue |
| GO:0070727 | cellular macromolecule localization | 0.033862353 |
| GO:0034613 | cellular protein localization | 0.03115501 |
| GO:1903827 | regulation of cellular protein localization | 0.034549815 |
| GO:0046907 | intracellular transport | 0.01168741 |
| GO:0051168 | nuclear export | 0.0122513 |
| GO:0051169 | nuclear transport | 2.56E-04 |
| GO:0006913 | nucleocytoplasmic transport | 0.014728889 |

**Table S4.**
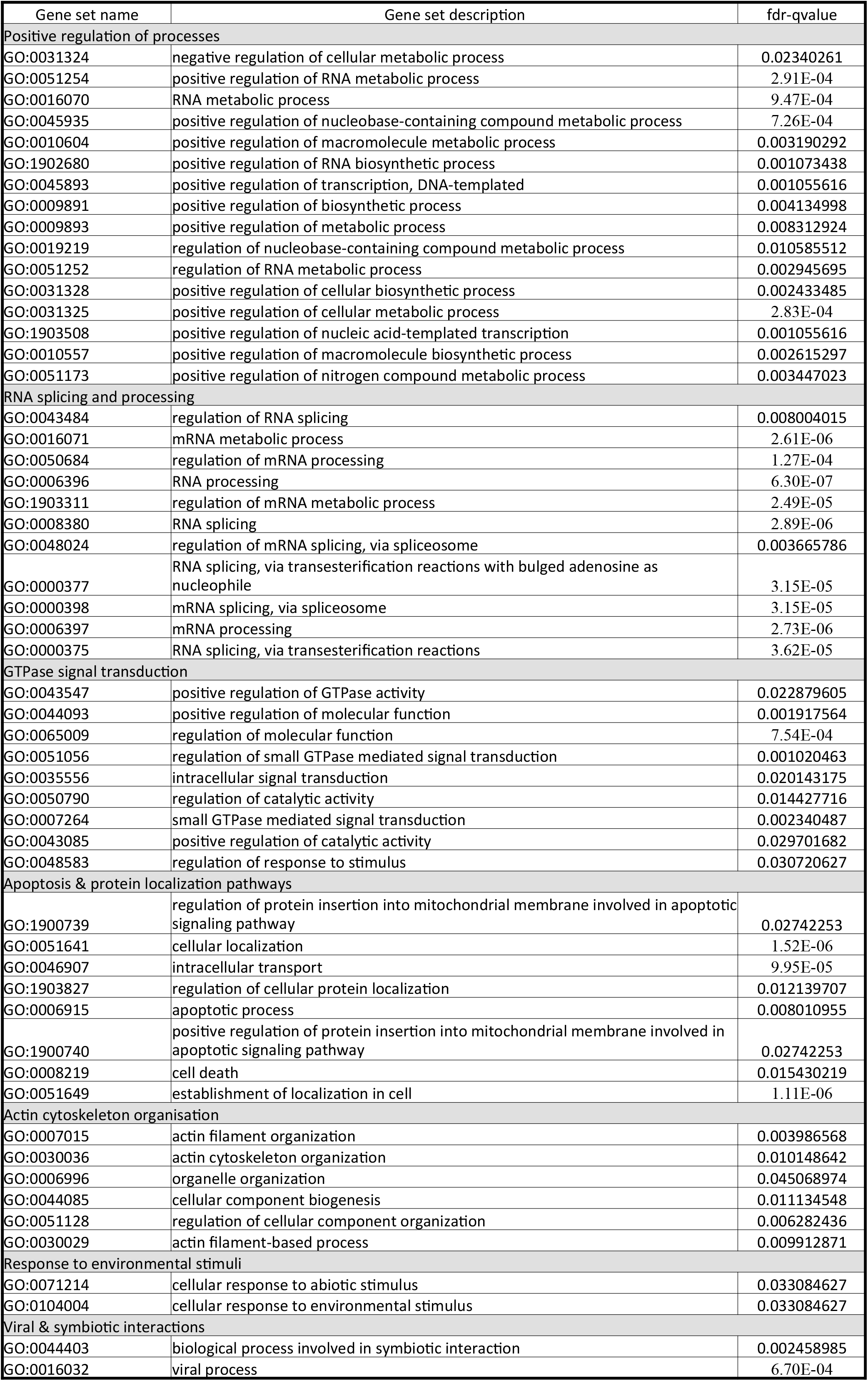
List of annota1ons of L-CK1.2 host substrates.

